# Phase leads between oscillatory visual stimuli induce a salience illusion

**DOI:** 10.1101/2022.10.31.514570

**Authors:** Benjamin J. Stauch, Eleni Psarou, Rasmus Roese, Pascal Fries

## Abstract

Neuronal populations often engage in oscillations, and corresponding theories and computational models propose that this affects their impact on other neurons. Specifically, when two neuronal populations oscillating at similar frequencies compete for impact, the phase-leading one might have an advantage by providing inputs to target neurons earlier. Here, we provide direct empirical support by driving visual neuronal responses with oscillating stimuli. We superimposed two orthogonal grating stimuli of temporally oscillating intensities, whose phase relations were precisely controlled. The leading stimulus was perceived as oscillating more intensely. This held for phase leads of a few milliseconds, and for each participant and almost every trial. Thus, we found a strong perceptual illusion directly predicted by theories on the functional role of oscillations for neuronal interactions.

## Introduction

Flexible cognitive functioning requires flexible modulation of neuronal communication. This is particularly evident for selective attention, where the attended stimulus is preferentially processed and perceived (Reynolds and Heeger, 2009). Such preferential processing might be implemented by oscillatory neuronal activities, together with mechanisms that give the attentionally selected neuronal activation a phase or time lead over competing activations (Fries, 2015; Palmigiano et al., 2017; Tiesinga and Sejnowski, 2010). The leading activation might selectively entrain higher-area neuronal groups, consisting of excitatory and inhibitory units, such that repeated cycles of the leading activation consistently arrive at phases of low network inhibition. By contrast, a lagging activation arrives at slightly later phases (and times) with high network inhibition and is thereby effectively suppressed. (Fries, 2015; Ni et al., 2016, Fig. 1A).

**Figure 1.**
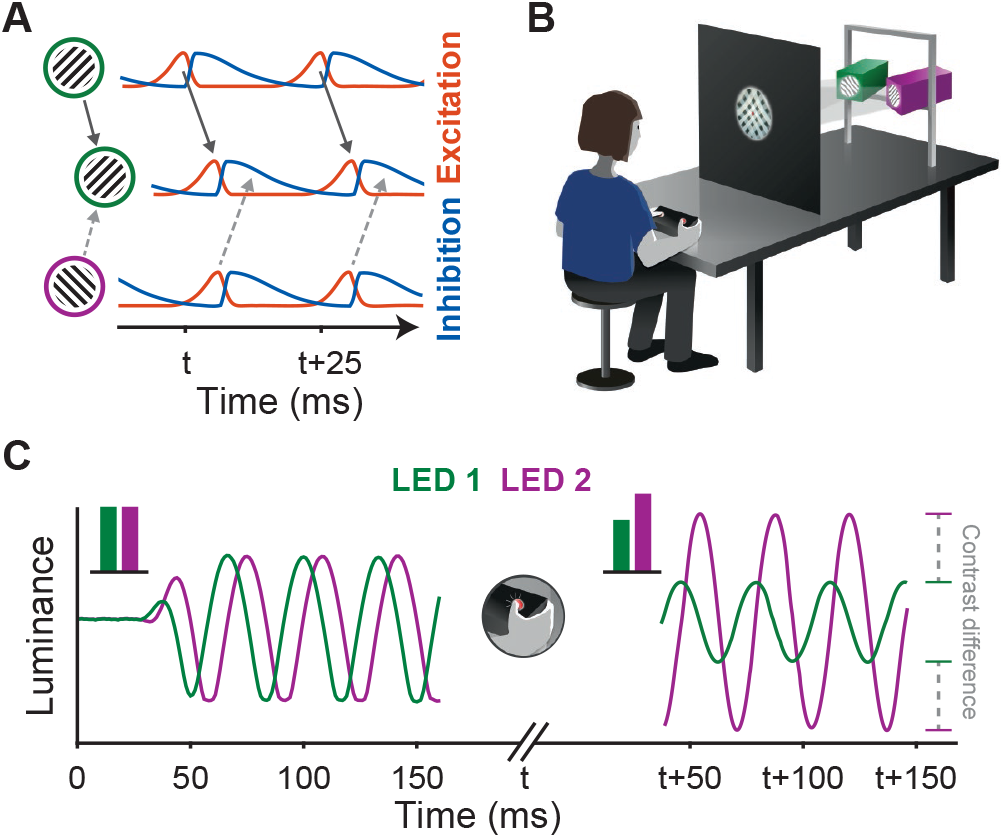
Experimental rationale and design. **(A)** Two presynaptic neuronal populations provide inputs to a common postsynaptic population. The presynaptic populations are driven by different stimuli (labeled green and purple), namely two gratings of different orientations. In each population, input-driven excitation (red) triggers network inhibition (blue), which inhibits the local network. The rhythm of the population driven by the green stimulus is phase-leading the rhythm of the population driven by the purple stimulus and has thereby entrained the postsynaptic population. Signals from the phase-leading presynaptic population now arrive at times of maximal excitability in the postsynaptic population, while signals from the phase-lagging population arrive at times of high network inhibition and are thereby blocked. Adapted from Fries (2015). **(B)** Experimental setup. Participants were seated in front of a projection screen, onto which two overlapping black-and-white gratings were projected from two individual LED projectors (labeled green and purple; same color scheme used for time courses in C). Participants controlled the relative stimulus oscillation amplitudes using a button box. **(C)** Luminances of the two stimuli, as measured on the projection screen. After stimulus oscillation onset, the stimulus projected from the left projector (green) was phase-leading the stimulus from the right projector (purple). At time t, the participant pressed the right button to increase the oscillation amplitude of the phase-lagging stimulus, thereby cancelling the perceptual effect of the phase-lead-induced illusion. The resulting difference in temporal contrast between the two stimuli was recorded. Note that stimulus onsets were tapered with half Hann tapers, such that both oscillation amplitudes rose smoothly at stimulus onset.

Indeed, when two neuronal groups engaging in biologically realistic oscillations are simulated in a computational model, the leading one has a substantially higher transfer entropy onto a joint target group (Palmigiano et al., 2017). Here, we empirically tested this proposal by providing two inputs with controlled phase relations. We hypothesized that the phase-leading stimulus benefits from enhanced processing and perception.

To drive neuronal activity in the early visual system with high temporal accuracy, we used oscillatory light stimulation (Bauer et al., 2012; Gulbinaite et al., 2019; Rager and Singer, 1998). Two superimposed oriented grating stimuli, whose orientations were offset by 90 degrees (to maximally separate the populations of driven neurons), were shown to 20 human participants (Fig. 1B). Their luminances were sinusoidally modulated at frequencies between 2 and 60 Hz. This frequency was fixed for a given trial, and identical for the two stimuli. Crucially, the phase relation between the two stimuli, and thereby their time relation, varied from trial to trial (Fig. 1C).

We hypothesized that the phase relation between the stimuli translates to a respective phase relation between the driven neuronal populations, and that the leading stimulus might be preferentially processed and perceived. On a given trial, one of the two stimuli was set to be phase leading or lagging, while keeping all other physical aspects constant, thereby isolating the effect of relative phase on perception. Thus, any perceptual difference would be a phase-relation induced visual illusion. Trials with a relative phase of zero were included as a reference condition.

Note that the absolute phase of both stimuli with regard to stimulus onset was chosen randomly for each trial, such that the order of which stimulus had an earlier onset was randomized. Note also that stimuli used in related previous studies often used pulses or square waves generated by typical monitors. Here, light-emitting diodes (LEDs; see Methods) were used to generate sinusoidal inputs of high temporal precision, which were likely more similar to visually induced oscillatory neuronal activity (Friedman-Hill et al., 2000; Kreiter and Singer, 1992; Livingstone, 1996).

## Results

The predicted illusion was indeed present. In a first experiment, participants were asked to write a free description, and all participants perceived the leading stimulus (here leading by 90°) as qualitatively more salient. Specifically, 55% of participants indicated that they saw only the leading stimulus flicker, 40% saw the leading stimulus flicker more strongly or faster, 15% described the leading stimulus as brighter, 10% as in front of the lagging stimulus, and 10% as having greater clarity or being more often in focus, whereby several participants reported multiple of these effects (Table S1).

To quantify the perceptual effect, a second experiment used the method of adjustment (again for 90° phase relations): Participants were again presented with two overlapping oscillating stimuli and were asked to modify the relative temporal contrast of the two stimuli until they perceived both of them as equal (Fig. 1C). The temporal contrast of a given stimulus, hereafter referred to simply as contrast, was quantified as the Michelson contrast between its maximal and minimal luminance in time (Max-Min / Max+Min). Participants modified the relative contrast between the two stimuli by pushing one of two buttons to simultaneously increase the contrast of one and decrease the contrast of the other stimulus, both in steps of 0.05. After participants had eliminated phase-lead-induced perceived contrast differences by adjusting physical contrast differences, the resulting difference [contrast(lagging stimulus) - contrast(leading stimulus)] quantifies the *perceptual contrast difference*. The perceptual contrast difference was bounded between zero (no physical difference) and one (one stimulus modulated with maximum amplitude, the other with zero amplitude). Participants took on average 12.3 seconds (*SD*_*participants*_ = 4.4 *s*) to register their choice.

For frequencies up to the known flicker fusion frequency (see below), a relative phase lead of 90° caused a sizable perceptual effect: To compensate for the enhanced salience of the leading stimulus, participants set the contrast of the lagging stimulus to a higher value. Figure 2A shows, for an example participant, the resulting perceptual contrast differences, separately for trials in which one or the other LED was leading. Figure 2B shows the average over both conditions and over all participants, revealing a phase-lead-induced perceptual contrast difference of 0.31 (*CI*_95%_ = [0.28, 0.34]). This corresponds to 31% of the maximal possible effect, and 62% of the average temporal contrast of the two stimuli. Figure 2C shows the reliability of the effect across presentations: On average, the leading stimulus was perceived as brighter (i.e., the perceptual contrast difference was positive) in 89% (*CI*_95%_ = [85%, 95%]) of all presentations below the threshold frequency of 37.4 Hz (*CI*_95%_ = [35.0 Hz, 38.6 Hz]).

**Figure 2.**
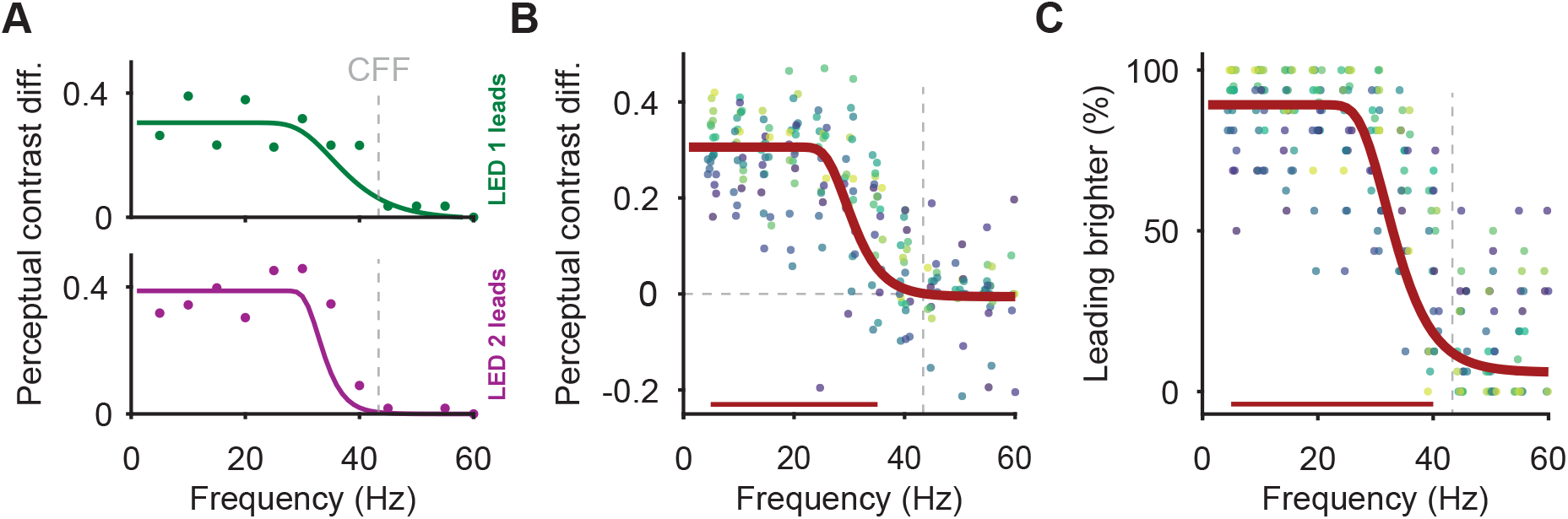
Effect strength and reliability over the perceivable frequency range. **(A)** Perceptual contrast difference (as defined in main text) per stimulus oscillation frequency in an example participant. Data are shown separately for trials in which one or the other LED was leading (upper and lower panel, respectively). Data were fit with four-parameter Gumbel functions (see Methods). The effect extended up to the known critical flicker fusion frequency (CFF Landis, 1954). **(B)** Average perceptual contrast difference for all participants (dot color indicating participant identity, dot x-position slightly jittered for visibility). Red line shows the fit averaged over participants. **(C)** Same as (B), but showing the percentage of trials with positive perceived contrast difference. Significance bar in (B) indicates significant effects above zero, in (C) above guess rate, both controlled for multiple comparisons across frequencies. For (B, C), *N* = 20 participants.

This effect persisted over the frequency range for which flicker is perceivable. The perceptual contrast difference was significantly above zero up to 35 Hz (Fig. 2B), and the reliability was significantly above chance up to 40 Hz (Fig. 2C). This is close to the flicker fusion frequency of 43 Hz (for human observers and for stimuli such as ours, i.e., central, 6 degrees visual angle diameter, medium luminance; Landis, 1954).

To probe the minimum phase/time lead that induced the illusion, we ran a third experiment: We again used the method of adjustment, now varying the relative phase/time lead between the two oscillating stimuli. Very small phase/time leads were sufficient to induce the perceptual effect: The reliability was above chance for phase leads of 22.5° or higher (Fig. 3A); this held for frequencies of 8 Hz, 16 Hz and 32 Hz, corresponding to minimal time leads of 7.81 ms, 3.91 ms and 1.95 ms (Fig. 3B). The effect increased with longer phase/time leads: maximal reliability occurred at 90°phase leads for all frequencies (see below for discussion). Qualitatively similar results were obtained for the perceptual contrast difference (Fig. 3D-E).

**Figure 3.**
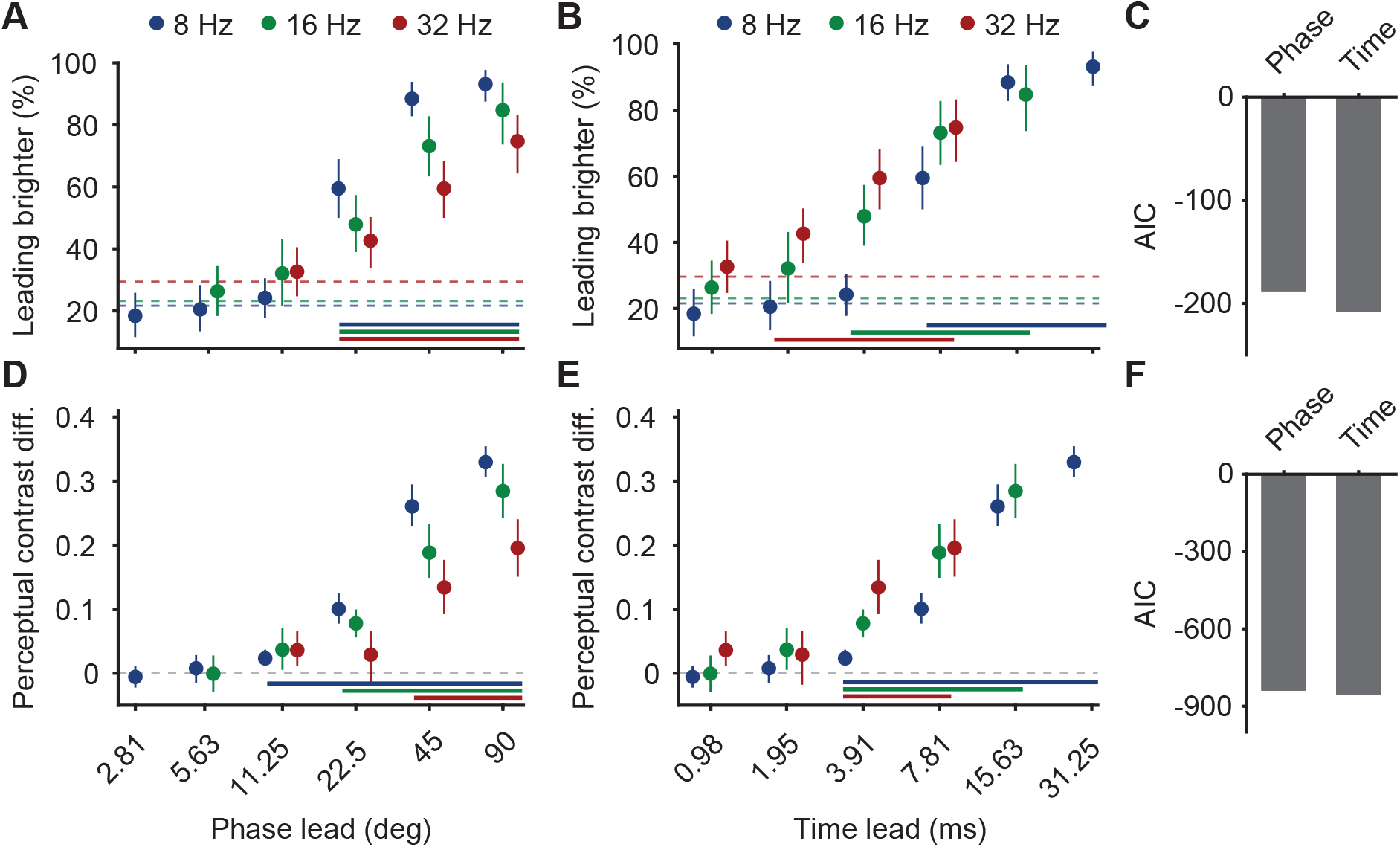
Effect strength and reliability for different phase and time leads. **(A)** Percentage of trials with positive perceptual contrast difference, averaged over all participants, separately for the different oscillations frequencies (legend on top) and the different phase leads. **(B)** Same data as (A), but after converting phase leads into corresponding time leads. **(C)** Model fit (as measured by AIC) of the data in A-B, fit with either phase-lead or time-lead predictors. **(D-F)** Same as (A-C), but replacing the percentage of trials with positive perceptual contrast difference by the average perceptual contrast difference. Dashed lines in (A, B) indicate per-frequency guess rates, calculated from the control condition with zero phase/time lead. Significance bars in (A, B) indicate significant effects above guess rate, in (D, E) above zero, each controlled for multiple comparisons across frequencies. Error bars in (A, B, D, E) indicate 95% bootstrap confidence intervals, *N* = 20 participants.

A given phase lead corresponds, for different frequencies, to different time leads, and vice versa. This allowed us to investigate whether the effect was better predicted by either phase or time leads. We therefore used either the phase leads or the time leads to predict either the effect strength (defined as perceptual contrast difference) or the effect reliability (defined as percentage of trials with positive perceptual contrast difference), using four separate hierarchical linear models. We then compared model fits: The time lead-based model showed a better fit than the phase lead-based model, both for effect reliability (*AIC*_*time*_ = −207.82, *AIC*_*phase*_ = −188.22, Fig. 3F) and for effect strength (*AIC*_*time*_ = −857.23, *AIC*_*phase*_ = −839.48, Fig. 3C).

## Discussion

In sum, human observers perceive the leading one of two oscillatory stimuli as more salient. The effect extends over the whole perceivable frequency range. Its strength increases with longer temporal offsets, yet it is already present for offsets as small as 1.95 ms. Its strength, as measured by the method of adjustment, corresponds to 31% of the maximal possible effect and 62% of the average temporal contrast of the two stimuli. Its reliability is 0.89, i.e., observers perceive it on almost every single trial. It was reported by each of 20 observers. Thus, it constitutes a reliable perceptual illusion, predicted on the basis of models suggesting that neuronal communication depends on neuronal phase relations (Fries, 2015; Palmigiano et al., 2017).

These models build on the observation that (rhythmic and non-rhythmic) fluctuations in cortical excitation are typically followed by corresponding fluctuations in inhibition, within 2-4 milliseconds (Atallah and Scanziani, 2009; Csicsvari et al., 2003; Hasenstaub et al., 2005; Vinck et al., 2013). This inhibition will suppress inputs that lag by as little as a few milliseconds, as observed in our experiments. While inhibition follows excitation within a few milliseconds, the net inhibition (i.e. inhibition minus excitation) peaks only at 90° after their joint peak, consistent over frequencies (Fig. S2A-B). Thus, the suppression of a lagging stimulus is expected to rise for phase lags up to around ≈ 90°, as we also confirmed.

This competitive stimulus interaction could take place at any visual processing stage that entails lateral inhibition and, crucially, that shows neuronal following responses to the stimulus oscillations. Neuronal flicker-following responses have been found from the retina up to visual cortical areas; they broadly decrease for higher areas, for foveal stimuli, and for higher frequencies (Rager and Singer, 1998; Van De Grind et al., 1973; Williams et al., 2004). These frequency limits of neuronal following responses might correspond to the known limits of flicker perception, and thereby to the limits of the effect observed here. There is indeed a broad correspondence between flicker perception and neuronal following responses (Van De Grind et al., 1973). Alternatively, neuronal following responses might exist for frequencies beyond the flicker fusion frequency, with potential consequences for the perception of isolated or competing stimuli that are more subtle. Yet, the robust illusion described here was intrinsically linked to flicker perception and the corresponding frequency range.

In summary, theories of phase-based stimulus selection and routing predict that phase-leading rhythmic neuronal inputs profit from a competitive advantage in further processing (Fries, 2015; Palmigiano et al., 2017). We show that a phase-leading stimulus is indeed perceived as being more salient. This establishes a novel visual illusion, directly based on theoretical predictions.

## ACKNOWLEDGEMENTS

We thank Michael Stephan for help in designing the LED driver chip. We thank Alina Peter, Patrick Jendritza, and Rasa Gulbinaite for insightful discussions on visual flickers and comments on a manuscript draft, and we thank Julia Trommershäuser for crucial advice regarding the task design.

## AUTHOR CONTRIBUTIONS

Conceptualization: BJS, EP, PF. Methodology: BJS. Investigation: BJS, EP, RR. Visualization: BJS. Funding acquisition: PF. Supervision: PF. Writing – original draft: BJS, PF. Writing – review & editing: BJS, EP, RR, PF.

## FINANCIAL INTERESTS

PF has a patent on thin-film electrodes (US20170181707A1) and is beneficiary of a respective license contract with Blackrock Microsystems LLC (Salt Lake City, UT), and he is member of the Advisory Board of CorTec GmbH (Freiburg, Germany). The authors declare that no further competing interests exist.

## Materials and Methods

### Participants

Participants were recruited via public advertisements, until 20 had completed all experiments (11 of them female). One participant completed only experiment 1; the remaining 20 participants completed all experiments. Participants were between 19 and 46 years old (median: 23.5 years), had never been diagnosed with any neurological or psychological disorder, did not regularly take medication except for contraceptives, and had normal vision or wore contact lenses. The study was approved by the ethics committee of the medical faculty of the Goethe University Frankfurt (Resolution 30/19).

### Stimulation setup

Participants sat in a darkened, sound-dampened room dimly illuminated by indirect DC LED lighting. They placed their head in a chin rest (including a forehead holder). A gray backprojection screen (Traverse 3D 140*140 cm, Stewart Filmscreen, Torrance, CA) was placed 56 cm from their eyes. Behind the screen, two identical custom-made slide projectors were placed. Each projector consisted of a cool white LED (LED ENGIN LZ1-00CW02, Osram Licht AG, Munich, Germany) mounted behind a circularly masked slide of a grating (either angled left or right by 45°), and a 25 mm diameter, 30 mm focal length achromatic lens (#27-267, Edmund Optics, Barrington, NJ).

Each projector produced a grating with a spatial frequency of 0.8 cycles/degrees visual angle, in a static circular aperture of 6.5 degrees visual angle diameter in the center of the projection screen. The projectors were mounted on a rail at equal height and angled inwards, so that their two projected grating patterns completely overlapped (Fig. 1B). Additionally, a red class-1 laser was used to backproject a fixation point of 2 mm diameter, which was placed in the middle of the joint aperture, where two dark grating stripes overlapped to form a black square.

To generate the LED-driving signal, we used a sound-to-LED pipeline implemented on a custom-made printed circuit board (PCB). Sinusoidal waveforms of per-trial varying frequency and phase were generated and played as audio output from Psychtoolbox running on a Windows PC (Kleiner et al., 2007), whereby the left and right audio channels were driving the left and right LED, respectively. A D/A converter (Chordette Gem, Chord Electronics Ltd., Kent, UK) was used as a USB soundcard. Its output was fed into the PCB consisting mainly of an operational amplifier (LM 358 PE3, Texas Instruments, Dallas, TX) and a DC offset (see Fig. S1) to transform the signal range into the operational range of the LEDs. Because the D/A converter output amplitude varies with output frequency, we measured the output to the LEDs for each frequency and adapted the per-frequency waveform amplitude output from Psychtoolbox until output to the LEDs was equalized across frequencies.

To shut off the LED-projected patterns between trials, we controlled a microcontroller board (Arduino Due, Arduino, Monza, Italy) to interrupt the LED circuits using a PhotoMOS relay (AQY211EH, Panasonic, Kadoma, Japan). The same Arduino also powered and controlled the fixation spot laser.

The illuminance reached by each of the two projectors for each possible temporal contrast setting was measured on the participant-facing side of the backprojection screen using a luxmeter (Luxcraft LX 1108, Conrad Electronic SE, Hirschau, Germany). When both stimuli were set to the same oscillation amplitude, each one oscillated between 442 and 1071 lx. The minimum and maximum oscillation ranges were 836-836 lx and 1-1435 lx, respectively.

The experimenter explained the tasks to participants and left the room after experiment 1. During all experiments, participants were monitored through an audio-video link.

### Paradigm

Throughout all experiments, participants were instructed to fixate the centrally presented fixation spot during the trials. Eye movements were recorded using an infrared eye-tracker (Eyelink 1000, SR Research, Kanata, Canada), and trials were aborted and restarted when participants looked away from the presented stimulus. Participants usually blinked several times during each trial. Participant responses were recorded using a response box (932, Current Designs, Philadelphia, PA).

Each trial proceeded as follows. Participants self-initiated the trial by fixating the fixation spot and pressing a trial-start button. Subsequently, the two gratings were shown statically for 200 ms. Then, their luminances began to oscillate, both with the same frequency and, for a given trial, with a fixed relative phase between them. The absolute phase (i.e. the starting phase along their oscillation) was randomly drawn for each trial to prevent stimulus-onset effects. Additionally, the oscillatory time courses were tapered at their beginning using the first half of a 100 ms long Hann taper (Fig. 1C) to prevent stimulus onset transients.

### Three experiments were conducted

First, participants were presented with gratings, whose luminances were both sinusoidally modulated at the same frequency (either 15 Hz or 35 Hz, randomly chosen for each trial). They were of equal luminance-oscillation amplitude and one of the two gratings was phase-leading the other one by 90° (randomly chosen for each trial). Participants self-initiated each trial, which lasted for 2.5 seconds. They were instructed to watch as many trials as they liked until they felt that they could fully express the perceptual differences between the two stimuli they saw. We attached small illustrations of the two separate, oriented grating stimuli to the upper left and upper right corner of the projection screen, respectively. We instructed participants to refer to the correspondingly oriented stimuli as “left” and “right” stimulus, when they described their perceptual experiences. Then, they were asked to write a sentence that would express the qualitative difference (Table S1). No prompts or examples of potential differences they might describe were given and no feedback or suggestions on the descriptions they formulated was provided.

Second, participants were presented with gratings, whose luminances were both sinusoidally modulated at the same frequency, now at frequencies of 5-60 Hz in 5 Hz steps, and with relative phases of 90°, 0° or −90°. Each combination of the 12 frequencies and 3 relative phases was shown 8 times per participant (for a total of 288 trials), with frequency and relative phase randomly permuted across trials. Participants were instructed to modulate the temporal contrasts of the two oscillating stimuli using button presses until they perceived both of them as equal. The temporal contrast of a given stimulus was quantified as the Michelson contrast between its maximal and minimal luminance in time (Max-Min / Max+Min). Participants pushed the left (right) button to simultaneously increase the contrast of the “left” (“right”) stimulus and decrease the contrast of the “right” (“left”) stimulus, both in steps of 0.05. Once they perceived both stimuli as oscillating equally, they pushed a trial-end button and thereby registered their choice and ended the trial. Participants had up to one minute to register their choice, but almost always responded earlier. If they did not register a choice during that time, the trial was aborted and restarted. No feedback was given.

The third experiment was identical to the second experiment, except that we varied the relative phase between the two stimulus oscillations. We presented participants with trials of sinusoidally modulated grating stimuli, both oscillating at either 8 Hz, 16 Hz or 32 Hz. Phase leads were chosen for each trial, such that they corresponded to time leads of ±[0 ms, 0.98 ms, 1.95 ms, 3.91 ms, 7.81 ms, 15.63 ms, 31.25 ms], and that phase leads did not exceed 90°. This resulted in phase leads of ±[0°, 2.8125°, 5.625°, 11.25°, 22.5°, 45°, 90°] for 8 Hz oscillations, ±[0, 5.625°, 11.25°, 22.5°, 45°, 90°] for 16 Hz oscillations, and ±[0, 11.25°, 22.5°, 45°, 90°] for 32 Hz oscillations. In total, 165 trials were run, such that each frequency-phase combination was shown five times.

### Analysis

For the first experiment, if participants provided their qualitative descriptions in German, they were translated into English using an online translation engine (DeepL Translate, DeepL SE, Cologne, Germany). All descriptions were classified into common description categories. The exact descriptions, as well as the translation and classification, are presented in Table S1.

For the method-of-adjustment experiments, we calculated two metrics, separately per participant and per stimulus condition (i.e. combination of frequency with phase lead, with the latter combining positive and negative phase leads of equal magnitude). The first metric, perceptual contrast difference (see main text), quantifies effect strength as the final contrast difference registered per trial, averaged over trials; values range between zero and one. The second metric quantifies effect reliability as the proportion of trials of each stimulus condition with positive perceptual contrast difference, i.e. the proportion of trials in which participants set the temporal contrast of the phase-lagging stimulus to a higher value than that of the phaseleading one; values range between 0% and 100%. To compute the guess rate of the reliability, we used the 0°phase lag condition, separately per frequency: We ran 1000 permutations in which we randomly assigned either of the two stimuli to be phase-leading or phase-lagging and computed the effect reliability as above. Statistical tests of the effect reliability in stimulus conditions with non-zero phase lags were computed against these guess rates.

For experiment two, we fit the resulting per-condition effect size and reliability values using inverted Gumbel functions with four free parameters (threshold, slope, upper asymptote, lower asymptote) using the psignifit 4 toolbox (Schütt et al., 2016). Then, the median parameter values were taken over participants to generate average fits of effect size and reliability, respectively, as a function of frequency. Confidence intervals of all parameters were computed using non-parametric percentile bootstraps. To analyze experiment three, we took the average effect size and reliability over all frequency-phase conditions and computed their confidence intervals, as above.

For both experiments, significance of effect strength against zero, and of reliabilities against guess rates, controlled for multiple comparisons, was computed using non-parametric t_max_-tests (Blair et al., 1994). Alpha was set to *α* = 0.05. Reported tests were one-sided, because experiment 1 had established the direction of effects.

## Supplementary information

**Table S1.** Verbal description of the induced percept in Exp. 1. Participants wrote their subjective perception of the illusion (Raw description), which was translated into English if needed (Translated description) and classified into one or two of the possible categories of percepts (Classification 1 & 2).

| Raw description | Translated description | Classification 1 | Classification 2 |
| --- | --- | --- | --- |
| Right one is more bright than left, while flickering. |  | Phaseleading pattern is brighter. |  |
| Einmal sieht die linkende [das linke] Gitter übersichtlicher aus als die rechtliche [das rechte] Gitter, und dann Gegenteil. | Once the left grid looks clearer than the right grid, and then opposite. | Phaseleading pattern has greater clarity. |  |
| Left is more fast while the right is stable. |  | Only phaseleading pattern flickers. |  |
| The right one looks constant and behind the left, while the left one is changing. I can see yellow border to the circle. |  | Only phaseleading pattern flickers. | Phaseleading pattern is in front. |
| Rechts flackert, während links still stand. Rechts starke Intensität des Flackerns. | Right flickers while left was still. Strong intensity of flickering on the right. | Only phaseleading pattern flickers. | Phaseleading pattern flickers more strongly. |
| L: pulsierend R: ruhig | L: pulsating R: steady | Only phaseleading pattern flickers. |  |
| Rechts flackert stärker als links. | Right flickers stronger than left. | Phaseleading pattern flickers more strongly. |  |
| The left moves faster than R. [Next trial:] R faster. [Next trial:] The R moves & L is stable. [Next trial:] L faster. [Next trial:] The L moves faster |  | Phaseleading pattern flickers faster. | Only phaseleading pattern flickers. |
| Trail 1: der linke Flicker stand still, während der rechte Flicker geflackert hat. Trail 2: der rechte Flicker stand still, während der linke Flicker sehr schnell geflackert hat. | Trial 1: the left flicker stood still while the right flicker flickered. Trial 2: the right flicker was still, while the left flicker flickered very fast. | Only phaseleading pattern flickers. |  |
| Der Fokus auf den beiden Muster wechselt innerhalb einer Präsentation. Anfangs war der Fokus auf dem linken Muster, [Next trial:] dann auf dem rechten [Next trial:] und dann wieder auf | The focus on the two patterns changes within one presentation. Initially the focus was on the left pattern, [Next trial:] then on the right [Next trial:] and then again on the left. The intervals of | The phaseleading pattern is more often in focus. |  |
| dem linken. Die Abstände des wechselnden Fokus waren gleich. | alternating focus were the same. |  |  |
| R hat geflackert L nicht | R flickered L not | Only phaseleading pattern flickers. |  |
| Links flickern langsamer am ersten Mal, am Gegenteil Rechts flickern schnellere. | Left flickers slower at the first time, on the contrary right flickers faster. | Phaseleading pattern flickers faster. |  |
| Links hat geflackert, Rechts blieb gleich | Left flickered, right stayed the same | Only phaseleading pattern flickers. |  |
| Der Rechte flickert schnell, der Linke gar nicht | The right one flickers quickly, the left one not at all | Only phaseleading pattern flickers. |  |
| Rechts wirkte statischer als links, links hat stärker geflackert als rechts | Right looked more static than left, left flickered stronger than right | Phaseleading pattern flickers more strongly. |  |
| Bei der letzten Präsentation flackerte das rechte Muster in nicht zu schnellen Geschwindigkeiten, während das linke sich nicht flackerte. | During the last presentation, the right pattern flickered at speeds that were not too fast, while the left pattern did not flicker. | Only phaseleading pattern flickers. |  |
| Der Linke flickert schneller als der Rechte. | The left one flickers faster than the right one. | Phaseleading pattern flickers faster. |  |
| Der rechte flickert schneller als der Linke | The right one flickers faster than the left one. | Phaseleading pattern flickers faster. |  |
| Links hell und intensiv flickernd, rechts dunkel und geringes flickern | Left bright and intense flicker, right dark and mild flicker | Phaseleading pattern is brighter. | Phaseleading pattern flickers more strongly. |
| Es sah so aus, als ob das linke Muster heller als das Rechte wäre und über das Rechte stehen würde. | It looked like the left pattern was brighter than the right one and stood over the right. | Phaseleading pattern is brighter. | Phaseleading pattern is in front. |
| Rechte bleibt stehen, unbeweglich und d[er] Linke flickert. | Right remains static, motionless, and the left flickers. | Only phaseleading pattern flickers. |  |

**Figure S1.**
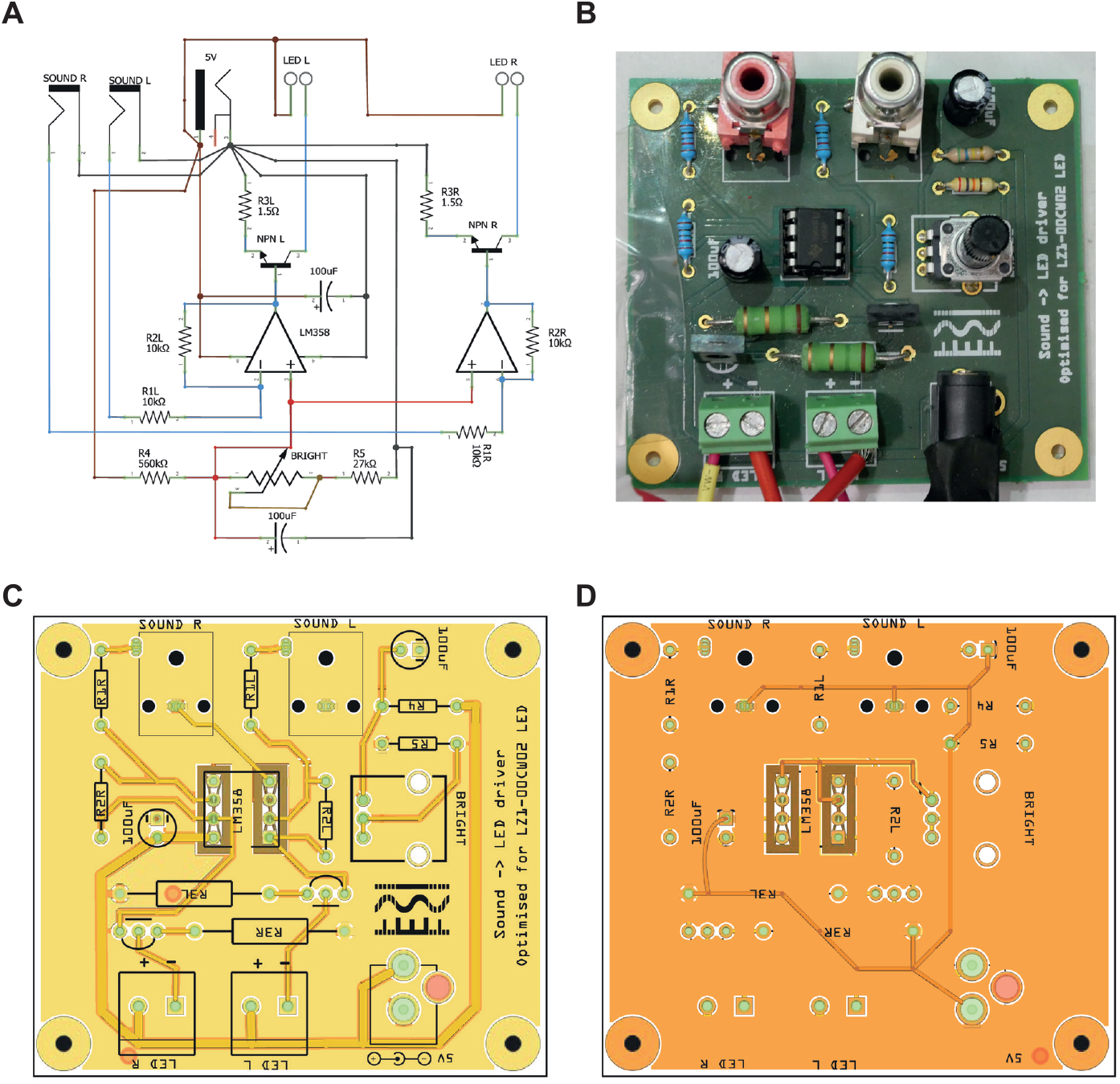
The LED driver board. **(A)** Circuit diagram of the LED driver board. Stereo sound input coming from the D/A converter was connected to SOUND R and SOUND L. Wires running to the two LEDs were connected to LED R and LED L. Between the LED driver board and the LEDs, a PhotoMOS relay was used to interrupt the LED power supply between trials. **(B)** A photograph of the assembled LED driver board. **(C)** PCB layout of the LED driver board, front. **(D)** PCB layout of the LED driver board, back.

**Figure S2.**
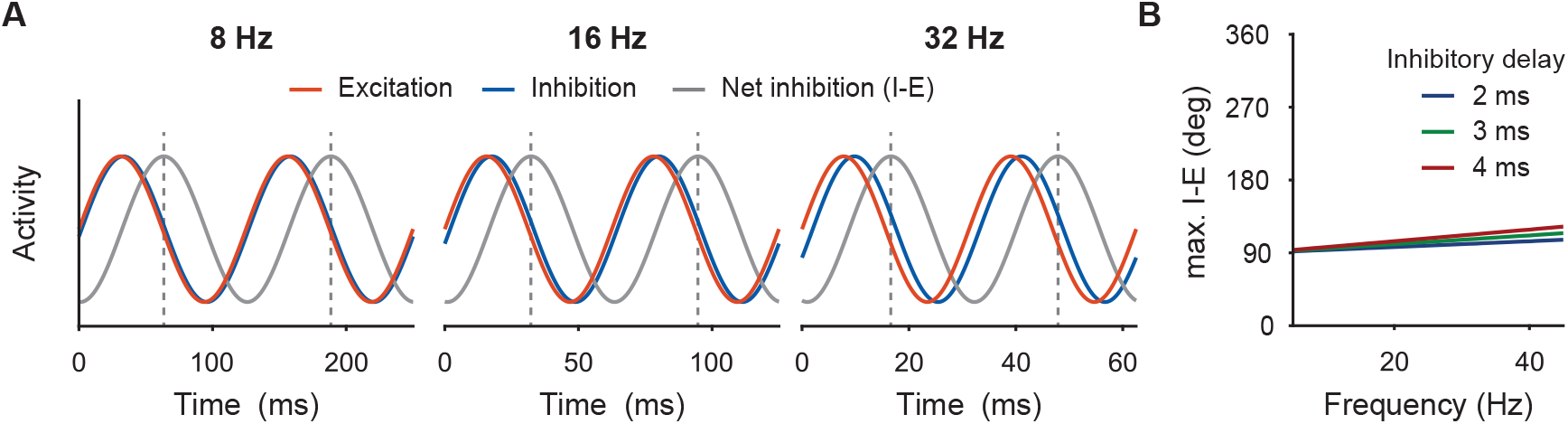
For short synaptic delays, 90° post-input shows strongest relative inhibition. **(A)** Schematic illustration of a sinusoidal excitation (red) followed, with a 2 ms delay, by sinusoidal inhibition (blue), separately for 8, 16, and 32 Hz. Gray lines show net inhibition, i.e., inhibition minus excitation, dotted gray lines show time points of peak net inhibition. Curves are scaled to their maxima; time axes are set to show two cycles per frequency. **(B)** Peak times of net inhibition, relative to the excitation peak, for the frequency range at which we found the effect and for different values of excitation-inhibition delays. As can be seen, all phases of peak net inhibition are at ≈90°or slightly afterwards.

